# HISTONE DEACETYLASE COMPLEX 1 modulates sepal length through the ethylene-ROS module

**DOI:** 10.1101/2025.03.27.645679

**Authors:** Dan Xiang, Dengying Qiu, Rui Zhang, Xi He, Shouling Xu, Ming Zhou, Lilan Hong

## Abstract

Organ size is precisely regulated during development to ensure proper function. The sepal must maintain an appropriate size throughout its growth to protect the developing flower bud until blooming. Despite its significance, the mechanisms regulating sepal size during its maturation remain poorly understood. Here we identify the HISTONE DEACETYLASE COMPLEX 1 (HDC1) as a positive regulator of sepal size during maturation in *Arabidopsis thaliana*. *hdc1* mutation prolongs sepal elongation due to delayed sepal maturation, thus hindering flowers from opening timely. Transcriptomic and proteomic analyses reveal that both plant hormone ethylene and reactive oxygen species (ROS) signaling are involved in the HDC1’s modulation of sepal size during maturation. Further genetic and chemical data show that during sepal maturation, HDC1 upregulates ethylene levels to promote ROS accumulation, thereby regulating the cessation of sepal growth. Collectively, our study revealed that HDC1 integrates ethylene and ROS signaling to regulate sepal maturation, providing new insights into the molecular mechanisms that govern organ size during maturation in plants.

## Introduction

Development is remarkably reproducible, generally producing organs with accurate size to ensure their proper physiological function, in an individual throughout their life (Chen *et al*., 2024a; Zhu *et al*., 2020). However, the behavior of cells that make up the organs is often variable and unpredictable, resulting in ambiguous contribution of cell size or number to organ size (Meyer and Roeder, 2014; Guo *et al*., 2010). There is persuasive evidence that plant and animal organs can sense their size and compensate it, through adjustment of their maturation time until the correct size has been attained despite the endogenous stochasticity at the molecular and cellular level as well as exogenous environmental fluctuations (Horiguchi and Tsukaya, 2011; Vogel 2013). Therefore, a deep understanding of the regulatory mechanisms underlying organ size control during maturation is critical and challenging in development biology.

Previous studies have revealed that plant hormones are involved in organ size during development. An *Arabidopsis thaliana* (*Arabidopsis*) MYB domain protein DEVELOPMENT RELATED MYB-LIKE1 (DRMY1) is required for both timing of sepal initiation and proper growth to maintain sepal size through affecting the spatiotemporal signaling patterns of auxin and cytokinin (Zhu *et al*., 2020). Further studies have revealed DRMY1 promotes the protein levels of ARABIDOPSIS RESPONSE REGULATOR 7 (ARR7) and ARABIDOPSIS HISTIDINE PHOSPHOTRANSFER PROTEIN 6 (AHP6) to suppress cytokinin signaling to control auxin patterning and sepal initiation (Kong *et al*., 2024). Besides, ethylene, one of plant hormones, has also been identified as an essential regulator of organ size. In cucumber, the S*hort Fruit 1* (*SF1*) gene encodes a cucurbit-specific RING-type E3 ligase that regulates the expression of *1-AMINOCYCLOPROPANE-1-CARBOXYLATE SYNTHASE 2* (*ACS2*, a rate-limiting enzyme gene in ethylene biosynthesis), controlling ethylene levels to influence cell division and fruit elongation (Xin *et al*., 2019). In *Arabidopsis*, the RING-H2 E3 ligase (RIE1) protein specifically interacts with and ubiquitinates AtACS7 to promote its degradation, thus affecting ethylene homeostasis to regulate leaf size (Tang *et al*., 2024). In *Solanum lycopersicum* (tomato), transcriptomic analysis of sepals from *MYB-Like Binding Protein 21* (*SlMBP21*) gene RNA interference transgenic plants, which exhibit longer sepals and improved fruit set, reveals that the differentially expressed genes were predominantly associated with ethylene and auxin signaling pathways (Li *et al*., 2017). Additionally, RNA sequencing analysis of sepal samples from *MADS-box* (*SlMADS1*) gene knock-out and *SlMADS1* overexpressing transgenic plants, which displayed altered sepal size, shows that genes related to ethylene, gibberellin, auxin, and cytokinin signaling are significantly affected (Xing *et al*., 2022).

Besides plant hormones, ROS also function as important signaling molecules that regulate nearly all aspects of growth and development in plants, including the control of organ size (Wang *et al*., 2024b). A MYB-like transcription factor, KUODA1 (KUA1) directly represses ROS generation to control leaf size (Lu *et al*., 2014). The increased accumulation of ROS level in *ftsh4-5* mutant, a loss-of-function mutant of mitochondrial i-AAA protease FtsH4, suppress sepal elongation (Hong *et al*., 2016). Recent study revealed that LATERAL ORGAN BOUNDARIES DOMAIN 11 (LBD11) and ROS modulate hypocotyl diameter in a negative feedback loop (Dang *et al*., 2023). However, whether and how ROS crosstalk with hormones to regulate organ size during its maturation still remains unclear.

Epigenetic modifications integrate various exogenous and endogenous signals to regulate growth and development. Histone deacetylation is one of important epigenetic modifications (Chen *et al*., 2024b; Deng *et al*., 2022) and HISTONE DEACETYLASE COMPLEX 1 (HDC1), a component of histone deacetylase complexes, plays a crucial role in regulating plant development through protein interactions in plants (Perrella *et al*., 2024). Studies have showed that HDC1 provides a potential link between histone H1 and histone-modifying complexes to regulate stress responses in *Arabidopsis* seedlings (Perrella *et al*., 2016, 2024). Additionally, HDC1 also interacts with HISTONE DEACETYLASE 6 (HDA6) to collaboratively regulate root system architecture remodeling under phosphate-deficient conditions in *Arabidopsis* (Xu *et al*., 2020; Feng *et al*., 2021; Li *et al*., 2022). However, the function of HDC1 in controlling organ size remains poorly understood, and further investigation is required to expand our knowledge.

The *Arabidopsis* sepals, the outermost floral organs, offer an ideal model system for investigating the mechanisms of organ size regulation for the following reasons: (1) An *Arabidopsis* flower has four sepals with the almost same size and an individual plant produces more than 50 flowers, allowing statistical assessments of organ size within a single organism, which generally cannot be done in animals. (2) Sepals are the outermost leaf-like floral organs, making them accessible for imaging during their maturation. (3) The size of a sepal is relatively insensitive to environmental effects, allowing us to focus on the intrinsic mechanisms that affect sepal size (Hong *et al*., 2016; Roeder, 2021).

In this study, through *Arabidopsis* mutant screening, we report that HDC1 regulates sepal size during maturation. In *hdc1* flowers, sepal maturation is significantly delayed, while the growth rate of the sepal remains unchanged, resulting in an elongated sepal. We show that HDC1 promotes 1-aminocyclopropane-1-carboxylic acid (ACC, the ethylene immediate precursor) accumulation and ACC synthase (ACS) enzyme activity, potentially leading to elevated ethylene levels to activate ROS accumulation, thereby promoting sepal maturation and terminating sepal growth. Our study reveal that that the HDC1 functions as a master regulator to integrate ethylene and ROS signaling to regulate sepal maturation, providing new insights into the molecular mechanisms that govern organ size during maturation in plants.

## Materials and Methods

### Plant materials and growth conditions

In this study, the *Arabidopsis* ecotype Colombia-0 (Col-0) is used as the wild-type (WT) plants. As described previously (Hong *et al*., 2016), mutants with increased sepal length were isolated from an M_2_ population of ethyl methane sulfonate-mutagenized Col-0. The *increased sepal length 1* (*isl1/hdc1-3*) mutated gene was identified through BSA-seq using the QTL-Seq pipeline (Huang *et al*., 2022b). The *isl1* (*hdc1-3*) mutation contains a G to A change at base 2703 of the coding sequence of HDC1, which generates a premature stop codon. The *isl1* (*hdc1-3*) mutation can be genotyped by amplifying with primers listed in data S5 at 55 °C annealing temperature, followed by sequencing. The *isl1* (*hdc1-3*) mutant was back-crossed to Col-0 three times to segregate unrelated mutations before further characterization. *isl1* (*hdc1-3*) was crossed with *hdc1-1* (allele with a T-DNA insertion in the first intron; GABI-Kat054G03) obtained from the *Arabidopsis* Biological Resource Center (ABRC; The Ohio State University) to test for allelism. All plants were grown in soil under a day-long cycle of 16 hours of light (∼100 µmol m^−2^ s^−1^) and 8 hours of dark at 22 °C in a growth chamber (Xu *et al*. 2024).

### Flower staging

Flowers at specific developmental stages were staged according to (Smyth *et al*., 1990) under a dissecting Leica microscope.

### Sepal length, width, and area measurements

The sepal length, width, and area measurements were performed according to previous description (Hong *et al*., 2016) with slight modification. The sepals dissected from stage 14 flowers were flattened between two slides and photographed on a black background using a dissecting Leica microscope mounted with a camera. Custom Python programs were used to extract each sepal’s contour from the sepal photos and to measure sepal’s length, width and area. The data were sorted, analyzed and plotted in Microsoft Excel or the statistical software GraphPad Prism 10.

### Cell area, cell number and cell length measurements

Mature sepal cell number and cell area measurements were performed according to previous description (Hong *et al*., 2016) with slight modification. Briefly, the sepals dissected from stage 14 flowers expressing pLH13 (35S::RCI2A-mCitrine) were imaged with a Nikon NIS-C2 confocal laser scanning microscope using a 20× objective. The stack images were processed in MorphoGraphX (Barbier *et al*., 2015) to segment individual cells and calculate cell area. Cell number was measured using the Fiji distribution of (Schindelin *et al*., 2012). Cell length was measured using ImageJ and MorphoGraphX.

### H_2_O_2_, ACC, 1-naphthaleneacetic acid (NAA), gibberellic acid (GA3) or aminoethoxyvinylglycine (AVG) treatment

The H_2_O_2_, ACC or AVG treatment was conducted as described previously (Li and Guo, 2018; Hong *et al*., 2016; Zhu *et al*., 2020) with slightly modifications. The primary inflorescences were dissected from indicated plants and then inserted into 1/2 MS medium-coated small petri dishes for at least 12 h for recovery. After recovery, the inflorescences were then transferred to 1/2 MS medium containing specific concentrations of H_2_O_2_, ACC or AVG, respectively and culture in a growth chamber for specific time.

For H_2_O_2_, NAA or GA3 treatment, inflorescences were dipped into 20mM or 40mM H_2_O_2_ solution, 5 µM NAA or 100 µM GA3 solution containing 0.05% (v/v) Silwet L-77, respectively. Distilled water with Silwet L-77 was used as a control. In this treatment, inflorescences were dipped once every day for 7 days.

### Detection and measurements of ROS

In situ detection of H_2_O_2_ and O_2_ ^• -^ were performed as described previously (Hong *et al*., 2016). For H_2_O_2_ detection, inflorescences were vacuum-infiltrated (three cycles of 5 min) with 0.1% (w/v) 3,3’-diaminobenzidine (DAB) in 10 mM sodium phosphate buffer (pH 4)/Tween-20 (0.05% v/v) and incubated in the dark (covered with aluminum foil) at room temperature overnight. For O_2_^• -^ detection, inflorescences were vacuum-infiltrated and incubated in 0.1% (w/v) nitroblue tetrazolium (NBT) in 50 mM sodium phosphate buffer (pH > 6.8)/ 0.05%Tween-20 (v/v) for 90 min at room temperature in dark. After reaching the optimal staining state, stained samples were removed from the staining solution and cleared by boiling in acetic acid:glycerol:ethanol (1:1:3, v/v/v) solution. The clearing solution was replaced once after the boiling. After clearing, samples (sometimes individual flowers were detached from the inflorescence if necessary) were photographed against a white background using a dissecting Leica microscope mounted with a camera.

### Ethylene signaling activity analysis by β-glucuronidase (GUS) staining

Ethylene signaling in young inflorescence was assayed using the *pEBS::GUS* reporter. GUS staining was performed as described (Sessions *et al*., 1999). The stained tissue was dehydrated and cleared with an ethanol series. GUS-stained seedlings were imaged with a digital camera mounted on a dissecting microscope.

### RNA isolation and quantitative RT-PCR (qRT-PCR) assay

Plant tissues were harvested and ground in liquid nitrogen. Total RNA was extracted using TRIzol reagent (Life Technologies) according to the manufacturer’s protocol. The first strand cDNA synthesis was obtained by HiScript II 1st Strand cDNA Synthesis Kit (Vazyme, Cat: R211-01) for gene clone, or by HiScript II QRT SuperMix for qPCR (+gDNA wiper) (Vazyme, Cat: R223-01) for qRT-PCR, following the protocol. For qRT-PCR, the cDNA samples were diluted and triplicate quantitative assays were performed with 2 µL of each cDNA dilution and 3 µL primers with the ChamQ Blue Universal SYBR qPCR Master Mix (Vazyme, Q312-02). The relative quantification method (- [[CT) was used to normalize quantitative variation between the replicates. The *UBIQUITIN 10* (*UBQ10*) gene was used as internal control. Three replicates were performed for each gene.

### Vector construction and plant transformation

To generate the *pHDC1*::*HDC1*-*mCitrine-3×Flag* vector for the complementation of *isl1* (*hdc1-3*) mutant, a genetic construct was designed to express the HDC1 protein fused to mCitrine and a 3*×*Flag tag under the control of the native *HDC1* promoter. The *HDC1* genomic sequence from the promoter (2557 bp upstream the start codon) to the sequence before the stop codon was amplified from Col-0 genomic DNA using PCR with Phanta HS Super-Fidelity DNA Polymerase (Vazyme, Cat: P502-d1). The PCR product was cloned into *pMOA33* vector (Barrell and Conner, 2006) at the the *Sac*I and *Spe*I sites with ClonExpress II One Step Cloning Kit (Vazyme, Cat: C112-02). The *p35S::mCitrine-RCI2A* vector was already available in the laboratory (Hong *et al*., 2016). All resultant vectors were verified by sequencing and transformed into the corresponding plants by *Agrobacterium*-mediated floral dipping. Seeds were selected on 1/2 MS medium with 50 µg/ml hygromycin. All primers used to construct the vectors are listed in Supplementary Table S1.

### RNA sequencing (RNA-seq) analysis

The sepals of WT and *hdc1-3* at stage 11 were dissected under a Leica stereo microscope and collected into 1.5 ml Eppendorf tubes floated in liquid nitrogen. Three biological replicates were collected, each replicate combining sepals from different individual plants. RNA extraction was done using a commercial kit following the manual. RNA integrity and concentration were assessed using an Agilent 2100 Bioanalyzer (Agilent Technologies), while the concentration was confirmed using the NanoDro 2000 (Thermo Fisher Scientific). Then the samples were sent to (Beijing Genomics Institute (BGI)) for sequencing on HiSeq 2500 (Illumina) platform in accordance with the manufacturer’s guidelines. Raw reads were subjected to cleaning and alignment to the TAIR10 *Arabidopsis* reference genome using HISAT2. Genes with a fold change greater than 2.0 and a *p* value < 0.05 were considered DEGs.

### HPLC-MS/MS analysis of ACC content and ACS activity

The sepals of WT and *hdc1-3* at stage 11 were dissected under a Leica stereo microscope and collected into 1.5 ml Eppendorf tubes floated in liquid nitrogen and each replicate combining sepals from different individual plants. Then the samples were sent to Nanjing Ruiyuan Biotechnology Co., Ltd. (China) for HPLC-MS/MS analysis. For ACC quantification, the plant material was cut, frozen in liquid nitrogen, and homogenized with deionized water at 4°C. After centrifugation, the supernatant was collected, concentrated, and filtered through a 0.22 µm filter. The samples were analyzed using a PE QSight 420 triple quadrupole LC-MS/MS system with optimized chromatographic conditions on a Poroshell 120 Aq-C18 column. Mass spectrometry was performed in positive ion mode using an ESI source for quantification. A standard curve was constructed with ACC standards, and the ACC content in the samples was quantified using the external standard method. The data were processed using PE QSight 420 software, and results were expressed in relative or absolute concentrations, followed by statistical analysis. For ACS activity analysis, the ACS crude enzyme solution was extracted from the samples using Tricine solution, followed by purification through a GE column to remove native ACC. S-adenosyl-L-methionine (SAM) was then used as the substrate to catalyze the production of ACC. The generated ACC content was measured using the PE QSight 420 triple quadrupole mass spectrometer (PerkinElmer, USA), and ACS activity was calculated based on the ACC concentration.

### Proteomic analysis

The sepals of WT and *hdc1-3* at stage 11 were dissected under a Leica stereo microscope and collected into 1.5 ml Eppendorf tubes floated in liquid nitrogen and each replicate combining sepals from different individual plants. Then the samples were sent to Shanghai Bioprofile Technology Co., Ltd. (China) for LC-MS/MS analysis in the positive-ion mode with an automated data dependent MS/MS analysis. Peptides from each sample were spiked with iRT standard peptides (Biognosys AG, Switzerland) and separated using reverse-phase high-performance liquid chromatography (RP-HPLC) on EASY-nLC system (Thermo Fisher Scientific, Bremen, Germany) with a column (75 µm × 150 mm; 2 µm ReproSil-Pur C18 beads, 120 Å, Dr. Maisch GmbH, Ammerbuch, Germany) at a flow rate of 300 nl/min. The DIA MS data were analyzed with Spectronaut 17 (Biognosys AG, Switzerland) (Bruderer *et al*., 2015; Barkovits *et al*., 2020). The MS data were analyzed for data interpretation and protein identification against the *Arabidopsis* database from Uniprot (downloaded on 10/17/2024, and including 136334 protein sequences), which is sourced from the protein database at https://www.uniprot.org/uniprotkb?query=arabidopsis&facets=model_organism%3A3702. Proteins with a fold change greater than 2.0 and a *p* value < 0.05 were considered DEPs. The overlapped DEGs and DEPs were obtained and used for associative analysis and GO enrichment using agriGO (http://systemsbiology.cau.edu.cn/agriGOv2/index.php).

### Combined RNA-seq and proteomic analysis

We used ID mapping (https://www.uniprot.org/id-mapping) to map protein identifiers to gene identifiers, enabling the identification of genes with differential expression in both the proteome and transcriptome. These results were then compared with the RNA-seq data to assess concordance and identify potential discrepancies between transcriptional and protein expression levels.

### Statistical analysis

All data were analyzed using GraphPad Prism 10 statistical software. The statistical significance among multiple datasets was determined using one-way analysis of variance (ANOVA) with least significant difference (LSD) tests; the statistical significance between two groups was determined using Student’s t-test.

## Results

### HDC1 modulates sepal length through regulating sepal maturation

To identify new components involved in regulating sepal size, we generated an ethyl methyl sulfonate-mutagenized population (over 5,000 mutants) from WT (Col-0) and then screened for mutants displaying altered sepal size. We found one mutant *increased sepal length 1* (*isl1*) displays longer sepals compared with WT (Fig. 1A). Further analysis revealed that compared with WT, *isl1* showed elongated flower (Fig. 1A, B) and increased sepal area, while the sepal width was not significantly changed in *isl1* (Supplementary Fig. S1A, B).

**Fig. 1.**
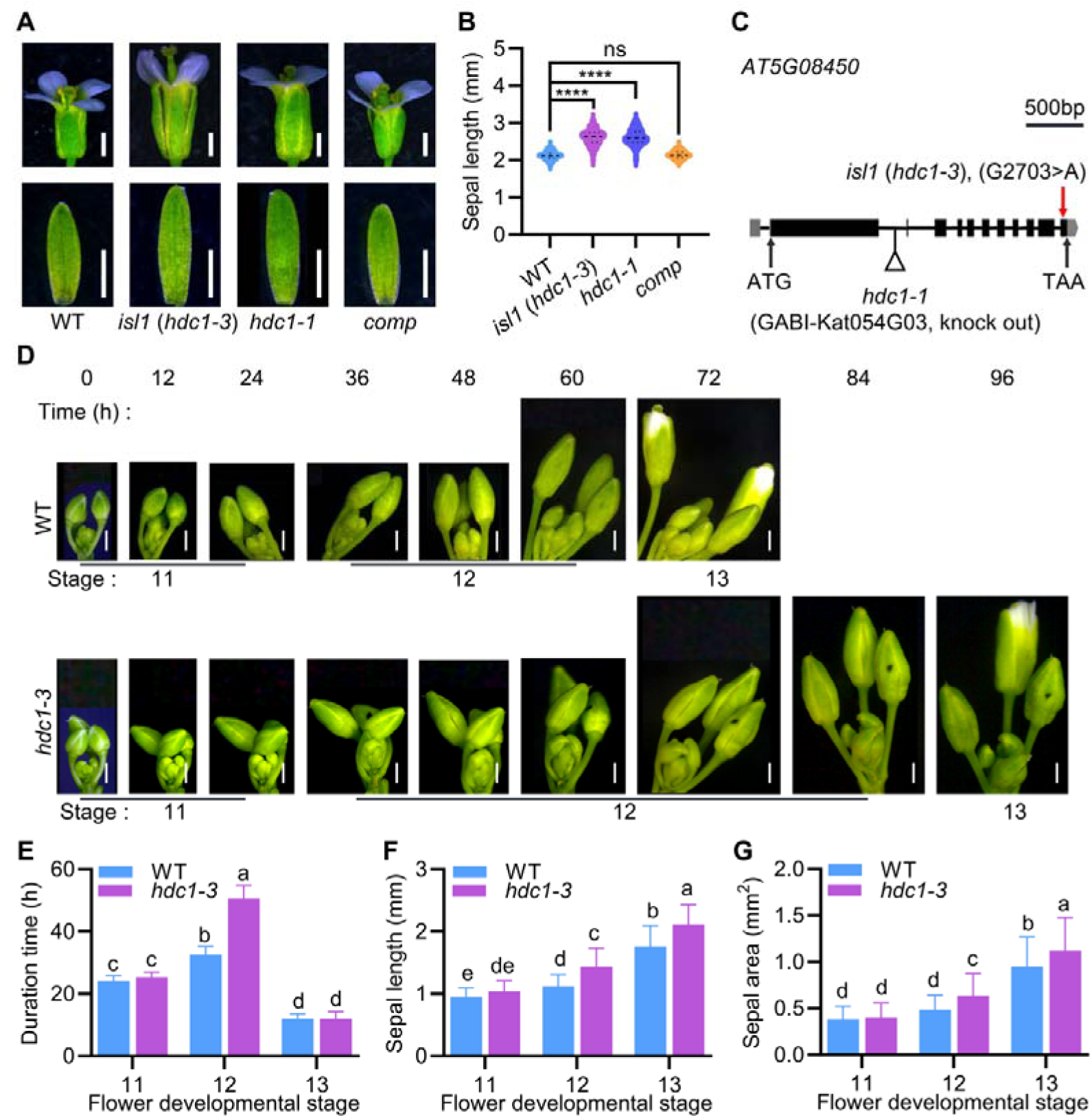
*HDC1* mutation leads to increased sepal length due to delayed sepal maturation. A: The mature flower and sepal morphology of wild-type (WT), *isl1* (*hdc1-3*), *hdc1-1* and the complemented line of *isl1* (*hdc1-3*) (*comp*). Bars = 1 mm. B: The lengths of WT, *isl1* (*hdc1-3*), *hdc1-1* and *comp* mature sepals. Data are presented using violin plots, n = 204 for WT, n = 183 for *isl1* (*hdc1-3*), n = 143 for *hdc1-1*, n = 130 for *comp*, ****p < 0.0001, ns for not significant, Student’s t-test. C: The gene model for the *HDC1* (*AT5G08450*). 5’UTR is shown in gray box, while 3’UTR is shown in gray pentagon. Exons are represented as black boxes, while lines represent introns. The *isl1* (*hdc1-3*) mutation consists of a point mutation of G to A, resulting in early termination, and the position of the T-DNA insertion in the *hdc1-1* mutant is indicated. D: The tracking of flower growth in WT and *hdc1-3* from stage 11 to stage 13. Bars = 1 mm. E: The duration time of stage 11 to stage 13 for WT and *hdc1-3* flowers. Data are means ± SD (n = 5). Different letters indicate significant differences (p < 0.01, ANOVA LSD test). F-G: The sepal lengths (F) and sepal areas (G) of WT and *hdc1-3* flowers at stage 11 to stage 13. Data are means ± SD (n = 5). Different letters indicate significant differences (p < 0.01, ANOVA LSD test).

To determine the candidate gene responsible for the phenotypes of *isl1*, we performed the bulked segregant analysis sequencing (BSA-seq) and identified one homozygous G-to-A nucleotide mutation at position 2,703 (G2,073A, counting from the start codon within genome sequence) in the gene *AT5G08450* which encodes a component of the histone-deacetylase complexes, HDC1 (Fig. 1C). This point mutation caused a premature stop codon, resulting in a dramatic decrease in *HDC1* transcript level and HDC1 protein accumulation (Supplementary Fig. S1C, D). Then we performed a complementation test by introducing a 6.1-kb WT genomic fragment (containing putative promoter region, entire ORF, and 3’ untranslated region) of *AT5G08450* into *isl1* and found that the transgenic plants displayed WT-like phenotypes, in terms of flower size, sepal length, and sepal area (Fig. 1A, B; Supplementary Fig. S1A, B). Additionally, we found that *hdc1-1*, a loss of function mutant (*GABI-Kat054G03*, Perrella *et al*., 2013) with a T-DNA insertion in the first intron of the *HDC1* gene, also showed *isl1-like* phenotypes (Fig. 1A-C). Moreover, our genetic analysis revealed that *isl1 hdc1-1* F_1_ hybrids exhibited *isl1-like* phenotypes (Supplementary Fig. S1E-H), indicating that *isl1* is allelic to *hdc1-1*. These results demonstrate that the mutation of *HDC1* promotes sepal elongation in *Arabidopsis*. Given that another T-DNA insertion allele, *hdc1-2* (*salk_043645*), was recently reported as broadly involved in environmental stress response (Feng *et al*., 2021), we renamed *isl1* as *hdc1-3*.

Then we systematically investigated the sepal length between WT and *hdc1-3* throughout their development. Results showed that there were no significant differences in bud length between WT and *hdc1-3* flower buds before stage 12, and the *hdc1-3* buds became gradually longer than WT buds from stage 12 to stage 16 (Supplementary Fig. S2A), which is consistent with the gradually increased expression levels of *HDC1* from early stages to stage 12 in sepals (Supplementary Fig. S2B). Further analysis revealed that compared with WT, the growth duration of flower buds at stage 12 was significantly increased in *hdc1-3* (Fig. 1D), Moreover, from stage 12 to stage 13, the sepal length, flower length, and sepal area were increased, but the sepal growth rate was not changed in *hdc1-3* compared with those of WT (Fig. 1D-G; Supplementary Fig. S2C-E). *Arabidopsis* flowers bloom at stage 13, when the sepals have reached full maturation. Our results show that *hdc1-3* mutants exhibit a delay in sepal maturation and an extended growth period, indicating that HDC1 regulates sepal length by affecting the timing of sepal maturation and growth.

### HDC1 regulates the proximal-distal cell length to modulate sepal length

To determine the cellular mechanisms of HDC1-mediated sepal elongation, we used a cell membrane fluorescence reporter, *mCitrine-RCI2A*, under the control of 35S promoter to investigate the cell size and number in *hdc1-3* sepals. Given that the epidermal cells in different regions of mature sepals show differential sizes, we compared epidermal cell sizes in the proximal, medial and distal regions between mature WT and *hdc1-3* sepals (Fig. 2A). Results showed that compared WT sepals, in *hdc1-3* sepals, the cell area and cell length in the proximal-distal direction were dramatically increased (Fig. 2B-D), while cell width (the dimension in the medial-lateral direction) was not changed significantly (Supplementary Fig. S3A). Moreover, the sepal cell number were modestly reduced (Fig. 2E). These results suggest that the increased sepal length of *hdc1-3* results from the enlarged cell size.

**Fig. 2.**
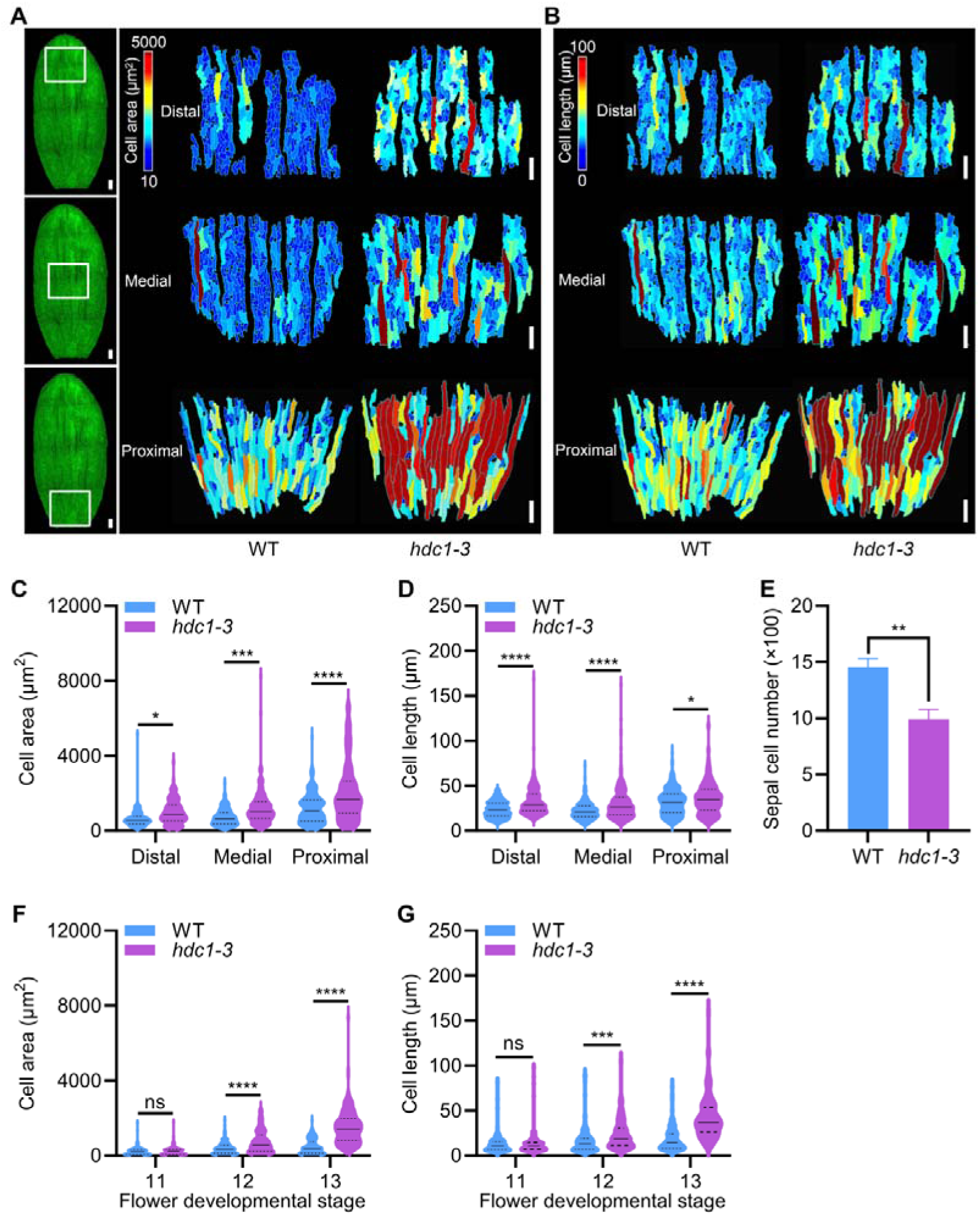
*HDC1* mutation increases sepal cell length in *Arabidopsis*. A: Heat maps show the areas of cells in the distal, medial and proximal regions of WT and *hdc1-3* sepals. Red and blue colors indicate large and small cells, respectively. Note that *hdc1-3* sepal exhibits a higher proportion of large cells and fewer small cells compared to WT sepal. On the left are maximum-intensity projection images for confocal stacks of WT sepals in which the epidermal cells are marked with a plasma membrane marker (green, *p35S::mCitrine-RCI2A*). The white boxes in the projection images show the distal, medial and proximal regions selected for cell geometry analysis. Bars = 100 μm. B: Heat maps show the lengths of cells in the distal, medial and proximal regions of WT and *hdc1-3* sepals. Red and blue colors indicate long and short cells, respectively. Note that *hdc1-3* sepal exhibit a higher proportion of longer cells and fewer shorter cells compared to WT sepal. Bars = 100 μm. C: Cell areas of WT and *hdc1-3* sepal epidermal cells from the distal, medial and proximal regions. Data are presented using violin plots, n ≥135, *p < 0.05, ***p < 0.001, ****p < 0.0001, Student’s t-test. D: Cell lengths in proximal-distal direction of WT and *hdc1-3* sepal epidermal cells from the distal, medial and proxima regions. Data are presented using violin plots, n ≥135, *p < 0.05, ****p < 0.0001, Student’s t-test. E: Epidermal cell numbers of WT and *hdc1-3* sepals. Data are means ± SD (n = 7). **p < 0.001, Student’s t-test. F: Areas of WT and *hdc1-3* sepal epidermal cells from stage 11 to stage 13. Data are presented using violin plots, n = 150, ****p < 0.0001, ns for not significant, Student’s t-test. G: Lengths in proximal-distal direction of WT and *hdc1-3* sepal epidermal cells from stage 11 to stage 13. Data are presented using violin plots, n = 150, ****p < 0.0001, ***p < 0.001, ns for not significant, Student’s t-test.

Then we investigated the cell sizes in WT and *hdc1-3* sepals from stage 11 to stage 13 to determine which stage is essential for the HDC1 coordinated sepal length regulation. Results showed that, compared to the WT sepals, the *hdc1-3* sepals exhibited a significant increase in cell area and proximal-distal cell length from stage 12 to stage 13, but not at stage 11 (Fig. 2F, G). In the meanwhile, the medial-lateral cell widths of WT and *hdc1-3* sepals were not significantly different from stage 11 to stage 13 (Supplementary Fig. S3B). These results suggest that HDC1 modulates sepal length through regulating the the proximal-distal cell length from stage 12 to stage 13 during sepal maturation.

### ROS are involved in HDC1-mediated sepal length regulation

To explore the molecular mechanisms underlying HDC1’s regulation of sepal length, we performed RNA-seq and proteomic analysis using WT and *hdc1-3* sepals at stage 11. We identified 1,238 differentially expressed genes (DEGs, log_2_ ≥ 1 or ≤ − 1; *p* ≤ 0.05), including 610 up-regulated genes and 628 down-regulated genes, and 430 differentially expressed proteins (DEPs, log_2_ ≥ 1 or ≤ - 1; *p* ≤ 0.05), including 257 up-regulated proteins and 173 down-regulated proteins, in *hdc1-3* versus WT (Fig. 3A, B; Supplementary Tables S2, S3). To clarify the pathways involved in HDC1’s function on the transcriptional and translational level, we processed a combined analysis and identified 57 DEG/DEPs overlapped in these transcriptomic and proteomic data (Fig. 3C, D; Supplementary Table S4). Gene Ontology (GO) analysis revealed that 51 HDC1-mediated DEG/DEPs were mainly and significantly enriched for functions including response to stress, catabolic process, redox homeostasis, response to hormone, defense response, metabolic process and cell wall biogenesis (Fig. 3D; Supplementary Table S5), indicating that HDC1 coordinates sepal maturation through regulating multiple intracellular regulatory pathways, especially redox homeostasis and hormone signaling.

Given the prominent enrichment of “redox homeostasis” in GO categories and the essential role of ROS in modulating redox homeostasis to regulate organ maturation (Hong *et al*., 2016). Then we examined endogenous O_2_^•-^ and H_2_O_2_ levels using NBT and DAB staining, respectively, in WT and *hdc1-3*. Results showed that compared with WT, the levels of O_2_^•-^ and H_2_O_2_ were all attenuated in *hdc1-3* inflorescences (Fig. 4A-C). Consistently, our qRT-PCR data further revealed that the expression of some ROS scavenging related genes were activated in *hdc1-3* sepals (Supplementary Fig. S4A), in line with the reduced ROS levels in *hdc1-3* sepals. Additionally, we examined the O_2_^•-^ and H_2_O_2_ levels in WT and *hdc1-3* sepals at different developmental stages to determine the stage starting to exhibit reduced ROS levels in *hdc1-3* sepals. Results showed that compared with WT sepals, *hdc1-3* sepals gradually exhibited the reduced O_2_^• -^ and H_2_O_2_ levels from stage 10 to stage 14 (Fig. 4D, E). All together, these results suggest that *HDC1* mutation affects ROS levels in sepals.

**Fig. 3.**
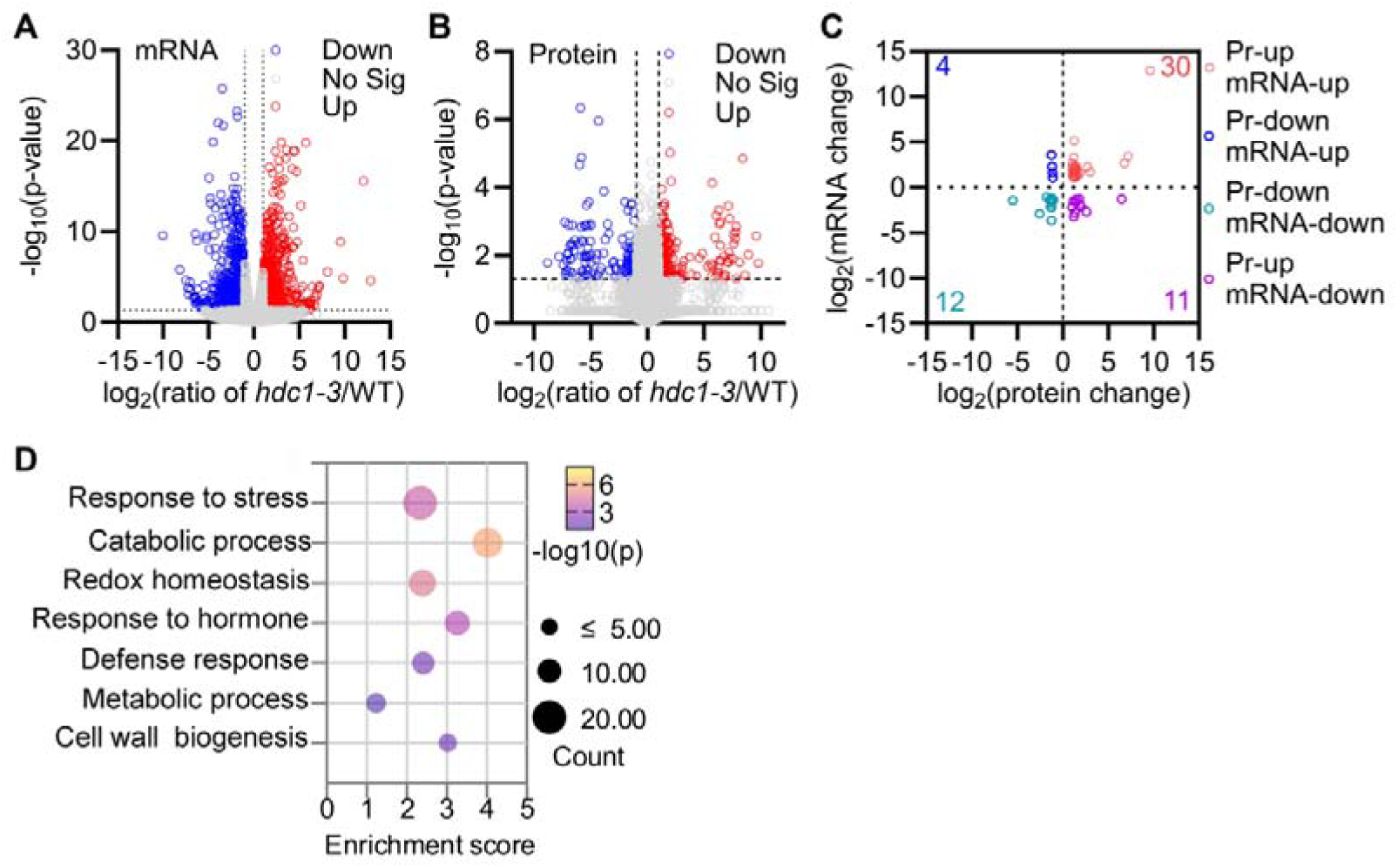
Transcriptomic and proteomic analysis of deferentially expressed genes and proteins of *hdc1-3* versus WT. A-B: Volcano plots of differentially expressed genes (A) and proteins (B) of *hdc1-3* versus WT (|log2 [fold change]| > 1, p < 0.05). C: Combined analysis of transcriptomic and proteomic data of *hdc1-3* versus WT (|log2 [fold change]| > 1, p < 0.05), Pr: protein. D: Gene ontology classification bubble plot of differentially expressed genes from transcriptomic and proteomic data in *hdc1-3* sepals versus WT sepals (|log2 [fold change]| > 1, p < 0.05). Larger bubbles indicate a greater number of genes associated with a specific GO term. Darker bubbles indicate smaller p-values, reflecting higher significance in the enrichment.

**Fig. 4.**
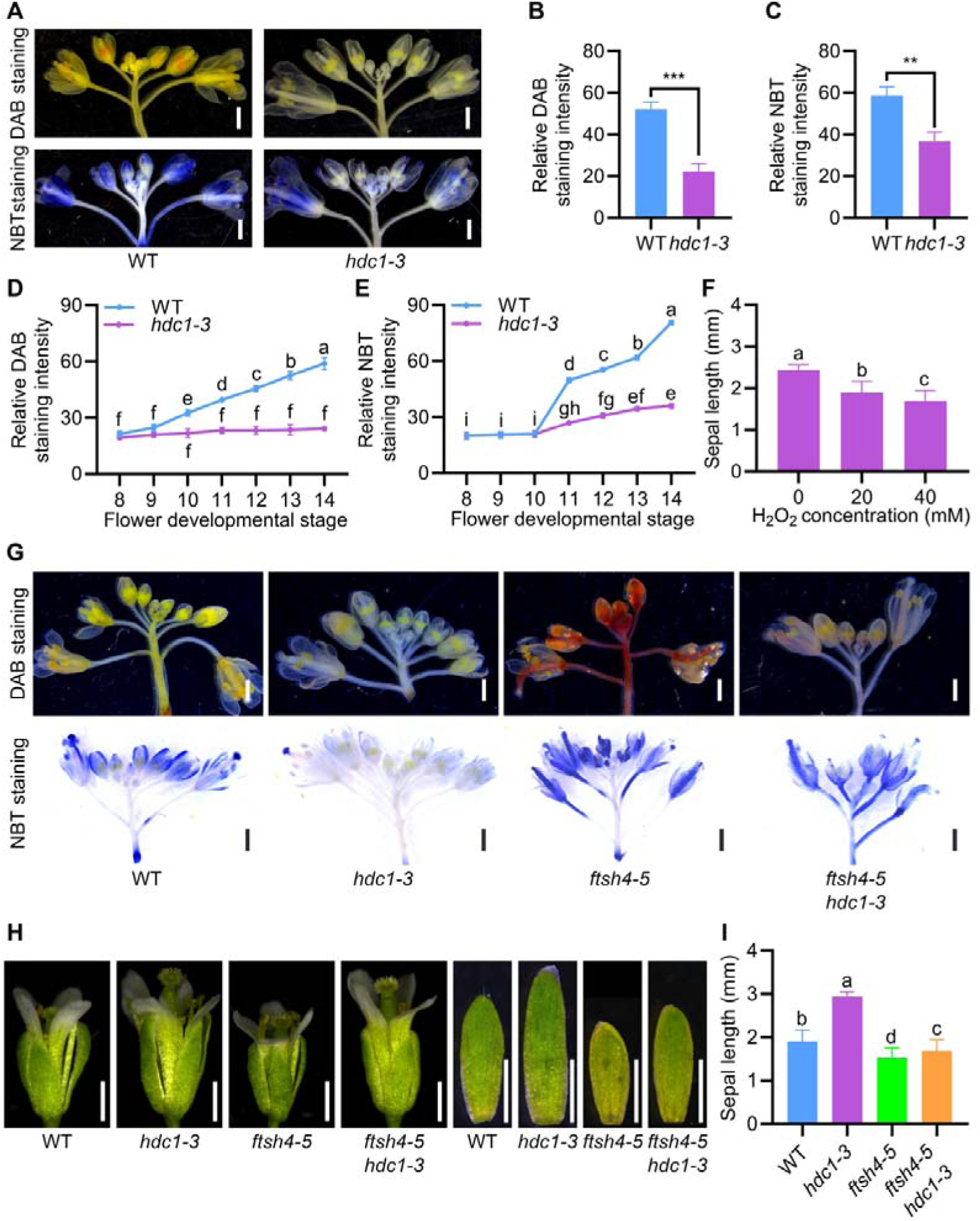
ROS signal is involved in HDC1-mediated sepal length regulation. A: The DAB staining and NBT staining of WT and *hdc1-3* inflorescences. Bars = 1 mm. B-C: Quantification of DAB staining (B) and NBT staining (C) intensities of WT and *hdc1-3* inflorescences. Data are means ± SD (n = 3), ***p < 0.001, **p < 0.01, Student’s t-test. D: Relative DAB staining intensities of WT and *hdc1-3* sepals at different developmental stages. Data are means ± SD (n = 3). Different letters indicate significant differences (p < 0.01, ANOVA LSD test). E: Relative NBT staining intensities of WT and *hdc1-3* sepals at different developmental stages. Data are means ± SD (n = 3). Different letters indicate significant differences (p < 0.01, ANOVA LSD test). F: The mature sepal lengths *hdc1-3* inflorescences treated with H_2_O_2_ of different concentrations. Data are means ± SD (n = 3). Different letters indicate significant differences (p < 0.01, ANOVA LSD test). G: The DAB staining and NBT staining of WT, *hdc1-3*, *ftsh4-5* and *ftsh4-5 hdc1-3* inflorescences. Bars = 1 mm. H: The flower and sepal morphology of WT, *hdc1-3*, *ftsh4-5* and *ftsh4-5 hdc1-3* at stage 13. Bars = 1 mm. I: The lengths of WT, *hdc1-3*, *ftsh4-5* and *ftsh4-5 hdc1-3* mature sepals. Data are means ± SD (n = 3). Different letters indicate significant differences (p < 0.01, ANOVA LSD test).

Then we investigated whether the lower ROS accumulation in *hdc1-3* is responsible for the delayed sepal maturation. To address this question, we increased the ROS levels in *hdc1*-3 sepals through H_2_O_2_ treatment, aiming to rescue the sepal phenotype. We treated WT and *hdc1*-3 flower buds at stage 8 with different concentrations of H_2_O_2_ for 7 days and then performed phenotypic analyses. Results showed that low concentrations of H_2_O_2_ (10 mM) treatment was not able to rescue sepal length of *hdc1-3* to WT level, but it was sufficient to reduce sepal length in WT (Supplementary Fig. S4B, C), showing a negative correlation between ROS levels and sepal length during maturation, in line with ROS’s role in promoting organ maturation (Hong *et al*., 2016). However, excessive application of exogenous H_2_O_2_ (40 mM through inflorescence dipping) dramatically suppressed the sepal length of *hdc1-3* (Fig. 4F, Supplementary Fig. S4D). Therefore, the lower ROS levels in *hdc1*-3 sepals hinder the maturation process promoted by ROS, and increasing ROS levels suppresses sepal elongation in *hdc1*-3 sepals, indicating that the longer sepals of *hdc1-3* result from its lower ROS accumulation. All in all, these results manifest that ROS are involved in the HDC1’s regulation of sepal maturation.

To further confirm the role of ROS in HDC1-mediated sepal mutation, we manipulated the ROS levels in *hdc1-3* mutant through genetics, by crossing *hdc1-3* with *ftsh4-5*, a precocious mutant that has reduced sepal length, due to elevated ROS levels (Hong *et al*., 2016). Phenotypic analysis showed that the reduced ROS levels, enlarged flowers, and elongated sepals of *hdc1-3* were all suppressed in *hdc1-3 ftsh4-5*, which showed similar phenotypes to *ftsh4-5* (Fig. 4G-I, Supplementary Fig. S4E, F), suggesting that elevated ROS levels brought by *ftsh4-5* mutation suppress the sepal elongation of *hdc1-3*.

The delayed sepal maturation and decreased ROS levels at later stages in *hdc1-3* sepals, together with our previous report on ROS working as signals promoting sepal maturation (Hong *et al*., 2016), further support the hypothesis that HDC1 regulates sepal mutation through modulating ROS levels in sepals.

### Ethylene is involved in HDC1-mediated sepal length regulation

Apart from “redox homeostasis”, the GO term “hormone responses” was also enriched significantly in the transcriptomic and proteomic data (Fig. 3D). To investigate the potential hormones involved in HDC1-mediated sepal length regulation, we accessed the effects of different hormone treatment on WT and *hdc1-3* sepal lengths. The results showed that the treatment of ethylene precursor ACC significantly reduced the sepal length of *hdc1-3* to WT level, but did not change the WT sepal length. Gibberellin treatment promoted sepal elongation in both WT and *hdc1-3.* NAA treatment slightly reduce sepal length in WT, while *hdc1-3* sepals were insensitive to NAA treatment (Supplementary Fig. S5A, B). Given ethylene plays essential roles in promoting organ maturation (Huang *et al*., 2022c), we chose ethylene to further explore its role in the HDC1-mediated sepal length regulation.

To verify ethylene’s roles on sepal length regulation, we first treated WT flower buds at stage 8 with different concentrations of ethylene biosynthesis inhibitor AVG or ACC for 7 days and then performed phenotypic analyses. Results showed that exogenous application of AVG dramatically increased sepal length and exogenous application of ACC dramatically decreased sepal length in WT (Fig. 5A, B) Additionally, we investigated whether endogenous ethylene levels also affect sepal length, using plants with different ethylene levels. We found that the sepals of *ACS6^DDD^*, which overproduces ethylene (Joo *et al*., 2008), showed decreased sepal length, and ACS-deficient mutant *acs octuple*, which exhibits reduced levels of ethylene (Mou *et al*., 2020), displayed elongated sepals (Fig. 5C, D). All together, these results suggest ethylene levels are correlated with sepal length negatively.

**Fig. 5.**
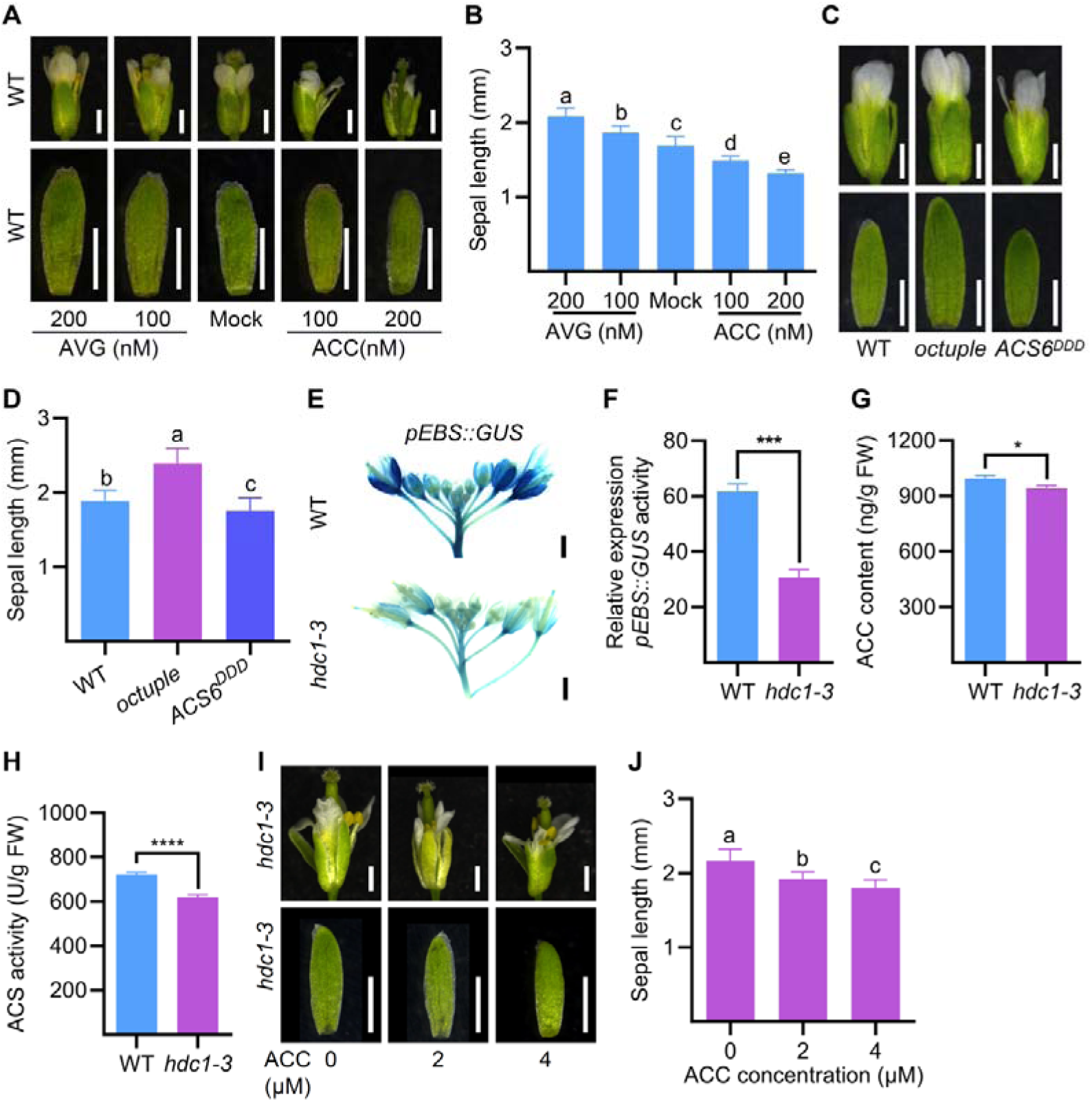
Ethylene signaling is involved in HDC1-mediated sepal length regulation. A: The mature flower and sepal morphology of WT after treated with AVG and ACC of indicated concentrations. Bars = 1 mm. B: The sepal lengths of WT flowers after treated with AVG and ACC of indicated concentrations. Data are means ± SD (n = 3). Different letters indicate significant differences (p < 0.01, ANOVA LSD test). C: The mature flower and sepal morphology of WT, *octuple*, and *ACS6^DDD^*. Bar =1 mm. D: Sepal lengths of WT, *octuple*, and *ACS6^DDD^*. Data are means ± SD (n = 3). Different letters indicate significant differences (p < 0.01, ANOVA LSD test). E: GUS staining of *pEBS::GUS* and *pEBS::GUS hdc1-3* inflorescences. Bars = 1 mm. F: GUS staining intensities in *pEBS::GUS* and *pEBS::GUS hdc1-3* inflorescences. Data are means ± SD (n = 3). ***p < 0.001, Student’s t-test. G-H:The ACC activities (G) and ACS contents (H) of WT and *hdc1-3* mature sepals. Data are means ± SD (n = 3). ****p < 0.0001, *p < 0.05, Student’s t-test. I: The mature flower and sepal morphology of *hdc1-3* after treated with and ACC of indicated concentrations. Bars = 1 mm. J: The sepal lengths of *hdc1-3* flowers after treated with ACC of indicated concentrations. Data are means ± SD (n = 3). Different letters indicate significant differences (p < 0.01, ANOVA LSD test).

Then we investigated the ethylene activity in WT and *hdc1-3* inflorescences using the ethylene reporter *pEBS:GUS* and found that the *pEBS:GUS* activity was reduced dramatically in *hdc1-3* compared with WT (Fig. 5E, F), suggesting ethylene activity is suppressed in *hdc1-3*. Additionally, we examined ACC (the ethylene immediate precursor) content in WT and *hdc1-3* and found that compared with WT, the ACC content was significantly reduced in *hdc1-3* mature sepals (Fig. 5G). Furthermore, we found that compared with WT, the activity ACS, the rate-limiting enzyme in ethylene biosynthesis, was also reduced dramatically in *hdc1-3* mature sepals (Fig. 5H). Besides, we treated *hdc1-3* flower buds at stage 8 with varying concentrations of ACC for 7 days and subsequently conducted phenotypic analyses. Results showed that exogenous application of ACC dramatically decreased sepal length in *hdc1-3* (Fig. 5I, J). These results suggest that *hdc1-3* sepals have compromised the ethylene activity, and *hdc1-3’*s longer sepal phenotype is related to its lower ethylene activity.

In summary, our results show that *hdc1* mutation leads to decreased ACS activity and ACC contents, and increasing ethylene levels rescues *hdc1* sepal phenotype. In addition, endogenous ethylene levels are negatively correlated with sepal length. Taken together, they suggest that HDC1 regulates sepal length through modulating ethylene levels, possibly through affecting ethylene biosynthesis.

### Ethylene induces ROS accumulation in sepals

ROS are now recognized as important regulators of plant developmental programs and also participate in various signaling cascades (Swanson & Gilroy, 2010; Singh *et al*., 2024). Under stress conditions, the production of ethylene is induced, which, in turn, stimulates ROS generation. These ROS act as signaling molecules, mediating the transduction of stress signals (Mittler *et al*., 2022). Having elucidated that both ethylene signaling and ROS were involved in HDC1-mediated sepal length regulation, we further examined whether ethylene also triggers ROS production in regulating sepal development. To address this question, we treated WT flower buds with different concentration of AVG or ACC at stage 8 and then examined the levels of O_2_^• -^ and H_2_O_2_. Results showed that compared with mock treatment, exogenous application of AVG significantly reduced the levels of O_2_^• -^ and H_2_O_2_, while exogenous application of ACC significantly increased the levels of O_2_^• -^ and H_2_O_2_ (Fig. 6A-C), indicating the positive correlation between ethylene levels and ROS levels in sepals. Moreover, we found that compared with WT, the levels of O_2_^• -^ and H_2_O_2_ were increased in the *ACS^DDD^*, whereas decreased in *acs octuple* mutant (Fig. 6D, E), further confirming the positive effect of ethylene on ROS accumulation in sepals.

**Fig. 6.**
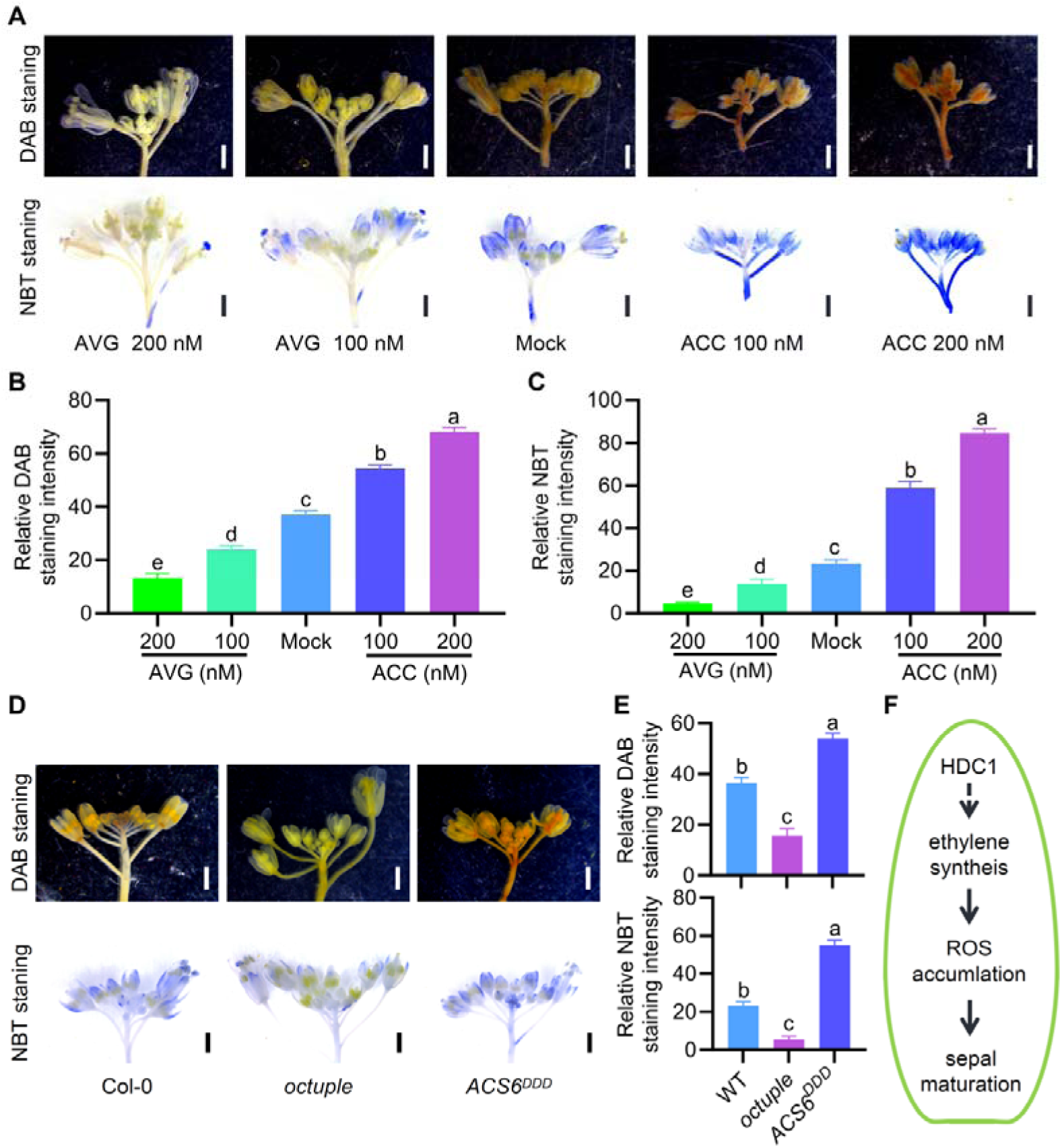
Ethylene induces ROS accumulation. A: The DAB and NBT staining of WT inflorescences after treated with AVG and ACC of indicated concentrations. Bars = 1 mm. B-C: The DAB staining intensities (B) and NBT staining intensities (C) of WT inflorescences after treated with AVG and ACC of indicated concentrations. Data are means ± SD (n = 3). Different letters indicate significant differences (p < 0.01, ANOVA LSD test). D: The DAB and NBT staining of WT, *octuple*, and *ACS6^DDD^* inflorescences. Bar = 1 mm. E: The DAB and NBT staining intensities of WT, *octuple*, and *ACS6^DDD^* inflorescences. Data are means ± SD (n = 3). Different letters indicate significant differences (p < 0.01, ANOVA LSD test). F: The proposed working model for HDC1-ethylene-ROS modulating sepal maturation in *Arabidopsis*. During the sepal development, HDC1 enhances ethylene signaling, which elevates ROS level to promote sepal maturation in *Arabidopsis*.

All in all, our analyses fit with a model that ethylene elevates ROS levels to promote sepal maturation, hence restricting sepal length.

## Discussion

Organs size controlling has captivated biologists for almost a century, yet the underlying mechanisms have just begun to be elucidated. In this study, using sepals as a model, we propose a novel mechanism of organ size regulation in which the HDC1-ethylene-ROS module plays a key role in maintaining proper sepal size during maturation in *Arabidopsis* (Fig. 6F).

HDC1 is an essential component of histone deacetylase complexes, primarily involved in regulating flowering time and environmental stress responses in plants (Ning *et al*., 2019; Xu *et al*., 2020; Feng *et al*., 2021; Zhou *et al*., 2021; Li *et al*., 2022; Liu *et al*., 2022; Patel *et al*., 2024; Kuang *et al* 2025). In this study we illustrate that HDC1 regulates sepal maturation. *hdc1-3* mutation delays sepal maturation and prolongs the time for sepal cell expansion. During sepal development, sepal cells expand asymmetrically with preferential growth along the proximal-distal axis (Hervieux, 2016; Hong *et al*., 2016). Therefore, *hdc1-3* mutants show increased length in sepal cells and sepals. The expression of *HDC1* is induced during sepal maturation (Fig. S2B), in line with its role in sepal maturation.

HDC1 promote sepal maturation through modulating ethylene levels. Our results demonstrate that *HDC1* mutation leads to decreased ethylene precursor contents and lower activity of major ethylene biosynthesis enzyme in sepals. Ethylene is one of the major hormones regulating organ maturation (Huang *et al*., 2022c). Ethylene biosynthesis genes *SlACS2* and *SlACS4* in tomato show important fruit-ripening functions (Upadhyay *et al*., 2023). During vegetative growth stage, ethylene also functions as a stimulator of cell proliferation under specific conditions (Zhu *et al*., 2011; Zhao *et al*., 2021). Thus the decreased ethylene levels in *hdc1-3* sepals might result in lower cell proliferation activity and fewer sepal epidermal cells, as well as delayed sepal maturation.

Numerous studies demonstrate that histone deacetylases are pivotal regulators of leaf senescence through their involvement in epigenetic control of gene expression (Liu *et al*., 2022; Huang *et al*., 2022a; Jin *et al*., 2024). Histone deacetylase complexes regulate transcription through both histone deacetylation-dependent and -independent mechanisms (Xu *et al*., 2023; Zhou *et al*., 2024; Patel *et al*., 2024). This process of gene expression regulation is controlled by histone acetyltransferases and deacetylases, which play crucial roles in cellular epigenetic regulation. Nevertheless, the detailed mechanisms through which HDC1 participates in epigenetic modification are complicated, and yet to be explored (Perrella *et al*., 2013, 2024; Feng *et al*., 2021). It is possible that HDC1 promotes sepal maturation through modulating the expression of genes involved in ethylene biosynthesis. However, our analysis showed that the expression levels of ethylene biosynthesis genes were variably changed in *hdc1-3*, with some genes upregulated, some genes downregulated, and a few genes not changed in expression (Supplementary Fig. S6). In light of the reduced ACS enzyme activity in *hdc1-3* (Fig. 5d), we speculate that HDC1 may not directly regulate the expression of *ACS* genes, although we can not rule out that the possibility that the *ACS* genes upregulated in *hdc1-3* inflorescences may be downstream targets of HDC1 regulation. Therefore, the precise molecular mechanisms through which HDC1 influences ethylene activity in organ maturation remain to be clarified, and will be an interesting topic for further investigation.

Apart from ethylene, the ROS signaling is also involved in HDC1-mediated sepal maturation. ROS signals have been shown to promote sepal maturation (Hong *et al*., 2016). Our findings reveal that ethylene induces ROS accumulation during organ maturation, providing new insights into the regulatory mechanisms of organ size at the maturation stage. Our research findings echo the observations that under stress conditions, ethylene production is induced, subsequently promoting the generation of ROS, which then function as signaling molecules to mediate following physiological changes (Yao *et al*., 2017; Jacobsen *et al*., 2021; Mittler *et al*., 2022; Wang *et al*., 2024a).

Overall, our study supports a model that during sepal maturation, HDC1 enhances ethylene biosynthesis and accumulates ethylene, which leads to elevated ROS level, thereby promoting sepal maturation and terminating sepal growth.

## Acknowledgements

We thank Prof. Hongwei Guo (Southern University of Science and Technology, China) for kindly providing the *ACS^DDD^* and *acs octuple* mutants, Prof. Ziqiang Zhu (Nanjing Normal University, China) for *pEBS::GUS* transgenic line, Prof. Jianli Yang (Zhejiang University, China) for *hdc1-1* mutant, Prof. Juan Xu (Zhejiang University, China) for providing Nikon NIS-C2 confocal platform. This work was supported by the National Natural Science Foundation of China (32270867 and 32070853), Startup Foundation for Hundred-Talent Program of Zhejiang University (to LLH and MZ), China Postdoctoral Science Foundation (2021M702855), and the Fundamental Research Funds for the Central Universities (226-2024-00102)

## Author contributions

DX and LLH designed the research; DX, DYQ, XH and SLX performed research; RZ and MZ analyzed the RNA-seq data; DX and LLH wrote the paper with the input of all authors.

